# Investigating the critical role of serine in antimicrobial peptide binding to lipopolysaccharide

**DOI:** 10.64898/2026.11.18.767556

**Authors:** Mohammadreza Niknam Hamidabad, Jyler Menard, R. A. Mansbach

## Abstract

Lipopolysaccharide (LPS), a major component of the Gram-negative bacterial outer membrane, is a potent endotoxin responsible for triggering severe host immune responses and sepsis. Understanding the atomic-level mechanisms governing how antimicrobial peptides (AMPs) recognize and neutralize LPS is essential for designing effective anti-endotoxin therapeutics. Here, we present an *in silico* investigation of the single-molecule interaction between *Escherichia coli* O111:B4 LPS and the cationic antimicrobial peptide GL13K, alongside two single-point mutants at position 9 (GL13KS9A and GL13KS9T), using all-atom molecular dynamics and umbrella sampling simulations. Consistent with experimental trends, mutating SER9 into alanine significantly alters peptide binding affinity to LPS. Surpisingly, removal of the polar character of serine leads to substantial changes in the interaction of the *N-terminus* with LPS via the loss of hydrogen bonding to stabilize a hook shape at the *C-terminal* end. In specific, while wild-type GL13K exhibits the strongest unbinding free-energy barrier, the S9A substitution changes the conformational landscape in such a way as to redirect the peptide’s N-terminus from lipid A toward the core oligosaccharide and substantially reduces the unbinding barrier, whereas S9T maintains an similar affinity. Our work demonstrates the effects of distal mutations even in peptides on their binding and lays important groundwork for the development of peptide-based LPS inhibitors.

## 1 Introduction

Lipopolysaccharide (LPS) is a vital component of the Gramnegative outer membrane (OM), which comprises about 75% of the OM ^1^. It is also one of the primary toxins secreted by many Gram-negative bacteria and is responsible for the over-stimulation of a host immune response that can lead to sepsis ^2^. Thus, it is a target of interest for drug design for two reasons. First, as a primary constituent of the OM, drug molecules that interact with it can localize to the surface of the bacteria, to either pass through into the cell to reach an internal target or to directly disrupt the membrane itself. Second, a drug molecule that interacts strongly with LPS may be able to neutralize its toxic effects on the host, either by accelerating host clearance or by disrupting its interactions with downstream signaling that leads to the damaging overexcitation of the immune system ^3,4^.

A typical LPS molecule is made up of three different components: (i) a hydrophobic moiety called lipid A, (ii) a hydrophilic core oligosaccharide, and (iii) a long hydrophilic polysaccharide repeating unit called the O-antigen ^5^. The lipid A moiety accounts for the vast majority of the endotoxic and immunostimulatory activity of LPS by binding to the Toll-like receptor 4/myeloid differentiation factor 2 (TLR4/MD-2) receptor complex and initiating downstream inflammatory signaling ^6^. It also anchors LPS in the bacterial outer membrane through its hydrophobicity and divalent cation-mediated electrostatic bridging of phosphate groups, resulting in tight packing ^7^. It is the most conserved structural moiety in LPS ^8^.

To combat Gram-negative bacterial infections, hosts rely on antimicrobial peptides (AMPs) as a first line of innate defense ^9^. Cationic AMPs are particularly effective because their positive charge drives initial electrostatic attraction to the negatively charged phosphate groups of lipid A/core moieties ^10^. Beyond host-derived peptides, a diverse class of broader LPS-binding molecules–ranging from serum proteins like lipopolysaccharide-binding protein (LBP) to engineered small-molecule antagonists– use these conserved electrostatic and hydrophobic features to neutralize endotoxins and disrupt outer membrane integrity ^11^. Peptide binders use these exact mechanisms. For instance, cyclic derivatives like polymyxin B gain a high affinity for lipid A phosphate groups to disrupt the outer membrane without systemic cell toxicity ^12^, while synthetic amphipathic peptides such as Pep19-2.5 reorder lipid A aggregate structures into biologically inactive multilamellar structures ^13^. Similarly, structural motifs derived from host-defense proteins such as human LL-37^14^ or 18-kDa cationic antimicrobial protein purified from rabbit leukocytes (CAP18) use amphipathic loops and basic residues to isolate endotoxins from binding to receptors and inhibit downstream inflammatory signaling ^15^, which offer a compelling therapeutic pathway. Because LPS neutralization mitigates toxicity without directly killing the bacteria, it minimizes the risk of driving antimicrobial resistance.

Among AMPs, the synthetic membrane-disrupting peptide GL13K, derived from human salivary proteins, exhibits potent bactericidal activity across multiple species of Gram-negative (and Gram-positive) bacteria, LPS-binding, and substantial reduction in endotoxin activity ^16^. Specifically, Abdolhosseini *et al*. ^16^ showed that GL13K administration significantly enhances survival in mice after LPS injection and identified residue 9 as critical for LPS binding. Furthermore, Ye *et al*. demonstrated that GL13K and its D-enantiomer form self-assembled amyloid-like nanofibrils upon interacting with bacterial surface components, including the *E. coli* O111:B4 LPS molecule ^17^. GL13K’s bactericidal activity has been linked to its ability to disrupt membranes through a proposed carpet/pore model that involves a substantial amount of *β* sheet content, although it does not appear to rely upon fibril formation ^**?**^.

Although interactions between AMPs and Gram-negative bacterial membranes have been investigated ^18–21^, the specific molecular-level interactions governing peptide binding to an isolated LPS molecule or LPS-micelle structures remain poorly understood, motivating a detailed investigation. Because LPS-binding of GL13K is known experimentally to be governed to a significant degree by a single residue, it is an excellent case study to shed light on the relevant molecular-level properties of peptide-LPS binding needed for LPS neutralization.

Previously, we showed how sequence and charge determine GL13K’s early aggregation and free energy landscape ^22^, then we expanded our study using constant-pH molecular dynamics (CpHMD) to explore single peptides and dimers across pH 7.0 to 12.5, revealing that while charge state regulates solution-phase behavior, the presence of a POPE/POPG membrane mimic at physiological pH 7.4 was the key factor for structural change ^23^. Specifically, proximity to anionic membrane headgroups stabilized a distinct *β*-hairpin conformation in the GL13K dimer that exposed residue 9 (SER09) to the solvent. This supports a model in which Gram negative inner membrane electrostatic interactions promote a structural transition required for antimicrobial activity. Combining this electrostatic-driven behavior with the role of Ser9 in LPS binding raises the key question of whether GL13K maintains its *β*-hairpin structure while interacting with LPS, the negatively charged outer membrane component of Gram-negative bacteria, how mutations at SER9 influence the peptide dynamics.

In this study, we employ molecular dynamics simulations to investigate the atomic-level interactions between an isolated *Escherichia coli* O111:B4 LPS molecule and the antimicrobial peptide GL13K alongside two variants, GL13KS9A and GL13KS9T. By mutating the serine at position 9, we investigate the specific role of serine in peptide conformation and spatial localization relative to the LPS molecule. Furthermore, to provide a rigorous quantitative analysis of the binding, we use umbrella sampling to estimate the binding affinities between the LPS molecule and each peptide variant.

## 2 Materials and methods

This section outlines the computational workflows and simulation protocols used to investigate the interactions between GL13K antimicrobial peptide variants and the *E. coli* O111:B4 lipopolysaccharide (LPS) system. Specifically, we describe the construction of the smooth LPS molecular model, the preparation of wild-type and variant peptide sequences, the parameters for both unbiased molecular dynamics and umbrella sampling simulations, and the tools chosen for analysis of the data.

### 2.1 Computational model of LPS molecule

Using the CHARMM-GUI LPS Modeler ^24^, we constructed a computational model of the *E. coli* O111:B4 LPS molecule, building upon previous findings that demonstrated GL13K interactions with this specific LPS type ^17^. Unlike laboratory strains such as *E. coli* DH5-*α* ^25^, which is a rough LPS phenotype lacking the O-antigen moiety, the complete O111:B4 smooth LPS architecture allows us to explicitly examine the O-antigen domain as an initial peptide binding region. We note that the *E. coli* O111:B4 label does not correspond to a fixed invariant LPS structure. Natural variations in both the core oligosaccharide and O-antigen (serotypes) composition have been reported, particularly within the core region ^26,27^. We chose to model our lipopolysaccharide (LPS) system based on *Escherichia coli* O111:B4 (Type 1 Lipid A with six acyl chains, an R3 core type, and four repeating O-antigen units) ^28,29^, as presented in Fig. 1-a. The model is structured into three regions and 31 residues as follows: the hydrophobic lipid A anchor (residue 1, ECLI), the inner core comprising two KDO residues (residues 2–3) and three heptose residues (residues 4–6), the outer core oligosaccharide region (residues 7–11), and the O-antigen domain spanning residues 12 to 31, which consists of four identical sugare repeating units (BGLC–AGAL–AGLC–ACOL–ACOL)

**Fig. 1.**
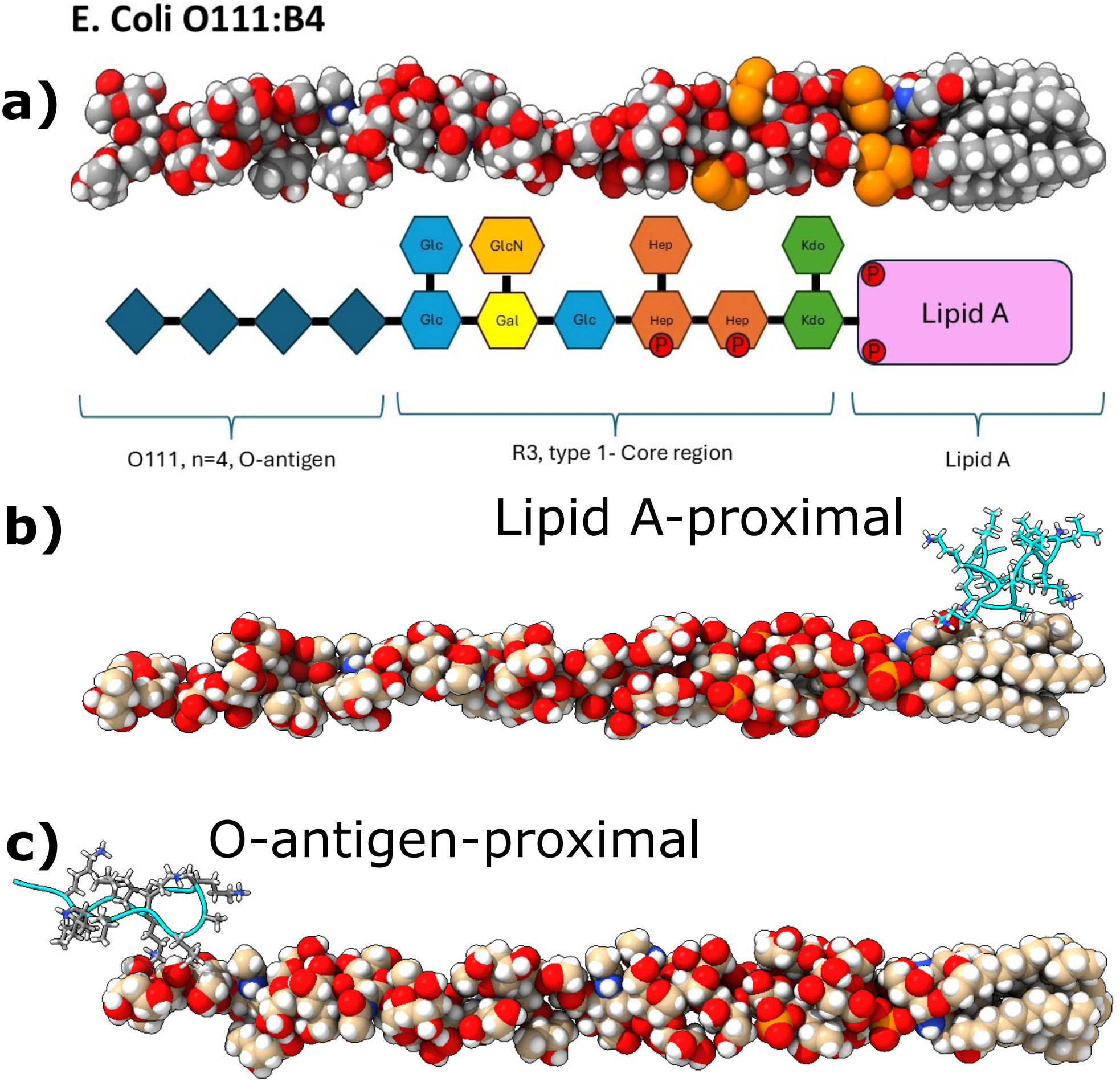
Computational model of the *E. coli* O111:B4 LPS system and initial peptide-LPS complexes. (a) Chemical structure of the *E. coli* O111:B4 LPS molecule annotated with schematic residue names. (b) Peptide-LPS complex starting structure with the peptide initialized at the lipid A region. (c) Peptide-LPS complex starting structure with the peptide initialized at the O-antigen moiety.

While previous studies have primarily investigated rough LPS chemotypes (such as those from *E. coli* DH5*α* mutants lacking the O-antigen) ^16,30^, modeling a smooth LPS chemotype is essential here because it allows us to investigate peptide interactions with the O-antigen layer in addition to the core and Lipid A regions. Furthermore, GL13K exhibits potent antimicrobial activity against *E. coli* O111:B4 in experimental studies ^17^. Using the full O-antigen repeating units provides a complete, physiologically relevant membrane representation to evaluate how these structural domains modulate peptide binding.

### 2.2 Initial peptide modeling

We prepared three peptide sequences: the wild-type GL13K and two variants, S9A (serine to alanine) and S9T (serine to threonine). We chose S9A for direct comparison to experimental work which used an alanine-scan to identify the ninth residue as the overwhelmingly relevant one for LPS binding, and S9T for a comparison with the most chemically similar amino acid, to identify the specificity of serine’s effects. The whole-peptide sequences are presented in Table 1. We introduced the mutations into the wildtype structure using PyMOL ^31^. The initial wild-type GL13K structure was selected from our previous study ^23^, in which the serine residue at position 9 is located at the tip of a hairpin-like turn, exposed to the environment. Consequently, the GL13KS9A and GL13KS9T variants share the same initial structure (Fig. 1b-c). We constructed all peptide variants with an amidated C-terminus and a protonated N-terminal glycine to match experimental conditions, as has been done before ^17,32,33^. With four lysine side chains and a charged N-terminus, each peptide carries a net formal charge of +5, which is what is expected at physiological pH conditions (pH 7.4).

**Table 1.** List of amino acid sequences studied in this report. The sequence differences from the original GL13K are highlighted in red.

| Peptide | Sequence |
|---|---|
| GL13K | *GKIIKLKASLKLL* |
| GL13K-S9A | *GKIIKLKAALKLL* |
| GL13K-S9T | *GKIIKLKATLKLL* |

### 2.3 Molecular dynamics simulations

We performed both unbiased MD simulations to characterize peptide regional bias, conformational changes, and unbiased peptide-LPS interactions and biased MD simulations to investigate the effects of residue substitution on the explicit peptide-LPS binding energy.

#### 2.3.1 Unbiased simulations

We first set up the initial starting structure of each peptide relative to the LPS molecule. Previous studies indicate that the lipid A and O-antigen moieties have distinct interactions with antimicrobial peptides (AMPs). The O-antigen acts as a hydrophilic barrier that can hinder direct peptide insertion toward the inner membrane, whereas the lipid A and core regions trap the AMP via electro-static interactions with anionic phosphate groups ^34^. Although manual placement near either region is possible, we employed a blind docking approach to ensure an unbiased and consistent initial starting structure across all three peptide variants. Initial binding poses of the peptides around the isolated LPS molecule were generated using the HPEPDOCK web server ^35^. Although HPEPDOCK is primarily designed for protein–peptide complexes, it was selected due to the absence of docking tools developed specifically for peptide–LPS interactions. According to the hypothesis generated from our previous work, we wanted to test whether a *β* hairpin conformation that exposes the ninth residue is particularly relevant for GL13K-LPS binding. To preserve that initial *β*-hairpin conformation at the beginning of simulation and ensure the residue 9 side chain remained exposed, we used global rigid docking using the pre-structured peptide rather than enabling flexible peptide docking. To account for potential limitations when applying a protein-peptide docking tool to a glycolipid system, we analyzed the top five results for each peptide and selected two representative starting complex structure: structure 1, in which the peptide interacts primarily with the lipid A/core region, and structure 2, in which it interacts with the O-antigen moiety. We expected structure 1 to be favorable under more realistic simulation conditions but employed both to ensure complete assessment of the role of starting structure in peptide-LPS interactions. In both systems, the peptide is placed approximately 1 nm from the LPS molecule, resulting in 6 different initial structures (2 initial structures per sequence).

We conducted all-atom molecular dynamics simulations of a single peptide interacting with LPS in solution using GROMACS ^36^ version 2021. We placed the LPS-peptide complex in a cubic simulation box with an edge length of approximately 15 nm, ensuring a minimum padding of 2 nm from each face to prevent periodic image interactions. We used the TIP3P water model ^37^ and filled the box. The system was neutralized using Na and Cl ions. Additional salt was not added to achieve physiological ionic concentration due to the fact that the primary objective was to characterize GL13K–LPS interactions while minimizing additional electrostatic screening. Divalent cations, such as Ca, were also not included, as their interactions with GL13K have not yet been systematically investigated computationally. Additionally, Ca is primarily known to stabilize LPS-LPS interactions ^7^, while in this system we focus on a single LPS molecule. We acknowledge that the absence of additional salt and divalent cations may influence electrostatic screening and the nature of peptide–LPS interactions. Therefore, the present simulations represent a simplified ionic environment rather than physiological conditions, allowing us to probe the effects of peptide chemistry alone in solution.

Electrostatic interactions were modeled using the particle-mesh Ewald (PME) method ^38^ with a real-space cutoff of 1.2 nm and a Fourier grid spacing of 0.14 nm. Van der Waals interactions were treated with Lennard-Jones potentials using a 1.2 nm cutoff. Temperature was maintained at 300 K using the velocityrescaling (v-rescale) thermostat ^39^ with a coupling time constant of 1 ps. Pressure was controlled at 1 bar using isotropic C-rescale pressure coupling ^40^ with a 5 ps period. Equations of motion were integrated using the leapfrog algorithm ^41^ with a 2 fs time step, and bond lengths involving hydrogen atoms were constrained using the LINCS algorithm ^42^ to fourth order with a single correction iteration. We simulated 5 x 2 starting structures = 10 independent replicates per peptide sequence (30 replicas in total) for 500 ns per replica and a total of 15 microseconds of simulation overall. All molecular dynamics simulations were performed using the Compute Canada (Digital Research Alliance of Canada) infrastructure on the Narval and Fir clusters. Computations were accelerated using NVIDIA A100 and H100 GPUs on Narval and Fir, respectively. Each GPU was assigned to each job with 4 CPU cores and 8 GB of memory on Narval, and shared 20 GB GPU configurations with the same CPU and memory on Fir.

#### 2.3.2 Biased simulations

To determine the binding free energy profiles (Potentials of Mean Force, PMF) between the peptides and the isolated LPS molecule, umbrella sampling ^43^ (US) simulations were performed. The reaction coordinate was defined as the distance between the center of mass (COM) of the peptide and the COM of the LPS molecule. Initial configurations for a series of windows were generated by pulling the peptide along the reaction coordinate from the bound state to the fully dissociated solvent region. We generated the initial structures by placing each peptide approximately 0.5 nm away from the Lipid A phosphate groups (P_A_), consistent with the peptide’s preference for Lipid A over the O-antigen. We then ran a 1 ns equilibration stage with position restraints applied to all LPS heavy atoms, allowing the peptide to relax near the interface. The final frame of this simulation was used as the starting configuration for the pulling phase. Harmonic restraints were applied to maintain the peptide within each window, and the resulting biased trajectories were combined using the Weighted Histogram Analysis Method ^44^ (`gmx wham` module) to construct the final PMF profiles and estimate binding free energies. The details and parameters related to the US method are described below.

The pulling step was performed to generate the biased configurations along a desired pathway. Starting from a bound state where the peptide center of mass (COM) was positioned approximately 0.5 nm away from the lipid A phosphate groups (*P*_*A*_), the peptide was pulled away from the LPS molecule. The pulling was constrained along the *y*-axis. Initially, 100-ns and 500-ns NVT and NPT equilibration were run, then the final frame was used as the initial structure for the pulling part. During the initial equilibration, position restraints were applied exclusively to the LPS molecule, allowing the peptide to interact freely. The pulling rate was set to 0.01 nm/ps (10 nm/ns) with a force constant *k* of 1000 kJ mol^*−*1^ nm^*−*2^. A series of 30 to 40 umbrella windows were generated, spaced at 0.1 to 0.2 nm intervals along the reaction coordinate. For each window, the system was run for 200 ps of NPT equilibration. Production runs for each window were carried out for 5 ns in the NPT ensemble. To ensure statistical reliability, this protocol was used for three independent replicas per peptide system.

### 2.4 Data Analysis

To comprehensively evaluate the conformational dynamics, folding behavior, intramolecular and intermolecular interactions of the GL13K antimicrobial peptide variants with lipopolysaccharide (LPS), we performed multi-step analyses. The following subsections provide a detailed description of these analyses: We start with residue-level contact propensities between peptide-peptide and peptide-LPS analysis, then we investigate the parameter shape of the peptides, and secondary structure propensities via DSSP assignments; later, we construct the conformational free energy landscapes projected onto collective variables and finally, the intramolecular and intermolecular hydrogen bonding calculations are presented.

#### 2.4.1 Contact Map Analysis

To monitor replica convergence and test whether there were effects of the starting structure, we calculated the COM distance between the entire peptide and the three distinct LPS regions, lipid A, the core oligosaccharide, and the O-antigen repeating units over the last 400 ns of the simulation using the MDTraj `compute_center_of_mass` module ^45^.

To quantify residue-level interactions between the antimicrobial peptides and the LPS molecule, pairwise distance matrices were computed using MDTraj `compute_center_of_mass` module and NumPy ^46^ `linalg.norm` module. For each trajectory, after fixing the periodic boundary conditions, the Center-of-mass (COM) coordinates were calculated for each individual peptide residue and each LPS residue within the lipid A and core oligosaccharide domains. Pairwise distances between the peptide and LPS residue COMs were evaluated over the final 400 ns of the production simulation. To ensure robust statistical sampling, data from the 10 independent replicas per peptide system were combined to calculate the overall mean distance and standard error of the mean (SEM) across all replicas.

#### 2.4.2 Shape Parameters

To quantify GL13K conformations, we calculate a variety of shape parameters using the MDTraj scientific package ^45^. We compute the radius of gyration (*R*_*g*_), acylindricity (*a*_cyl_), and asphericity (*a*_sph_), which are all quantities derived from the principal moments 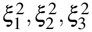 of the gyration tensor with the third component defined as the largest 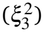. Additionally, we calculate the end-to-end distance of the peptide (*r*_*e*2*e*_) as the distance between *α* carbon of the first and the last residues. The radius of gyration is defined as

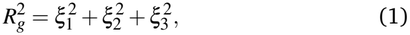

and is a measure of the overall compactness (low *R*_*g*_) or extendedness (high *R*_*g*_) of the protein.

Asphericity is calculated as

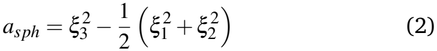

and shows how much the shape of a peptide deviates from a perfect sphere. A value of zero indicates a perfect sphere 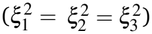, while the maximum value corresponds to 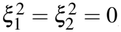, which results in 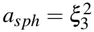. Under these circumstances, this leads to 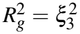, which indicates a perfect one dimensional line (thin rigid rod) along the peptide’s major principal axis.

Acylindricity, the analogous measure of deviation from a perfect cylinder, is calculated as

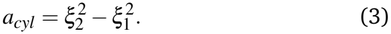

A value of zero indicates a perfect cylinder. We can describe the acylindricity as measuring the asymmetry of the mass distribution in the plane perpendicular to the main axis.

End-to-end distance is calculated as

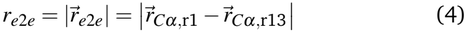

which measures the straight-line distance between the two terminal *C*_*α*_ atoms. Unlike the radius of gyration, this does not quantify the overall shape of the protein; a small *r*_*e*2*e*_ implies proximity of the termini but does not distinguish between a flat hairpin and a rounded loop. In combination with the other shape parameters, it gives a solid overall summary of the conformational shape, distinguishing between folded over versions of similar shapes.

#### 2.4.3 Secondary Structure Analysis

To quantify the local conformation of GL13K and its variants, we evaluated the per-residue secondary structure propensities across the 10 replicas of each system. Secondary structure assignments were determined using the DSSP algorithm as implemented in MDTraj ^45^. For each residue *R* (*R* ∈ {1, 2, …, 13}) and secondary structure type *S*, the ensemble population percentage *P*_*R,S*_ was calculated as:

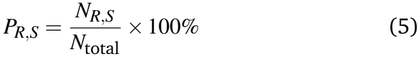

where *N*_*R,S*_ is the number of frames (aggregated across all independent production replicas) in which residue *R* was assigned to secondary structure state *S*, and *N*_total_ is the total number of analyzed trajectory frames, which is the final 400 ns of production trajectories from 10 independent replicas. Eight standard DSSP structural categories were evaluated: *α*-helix, isolated *β*-bridge, extended *β*-ladder, 3_10_-helix, *π*-helix, hydrogen-bonded turn, bend, and random coil/loop.

#### 2.4.4 Free Energy Surfaces

To characterize the conformational landscape of the peptides during interaction with the isolated LPS molecule, free energy surfaces (FES) were constructed as a function of two collective variables: the radius of gyration (*R*_*g*_) and the end-to-end distance (*r*_*e*2*e*_). While we could have automatically extracted collective variables using dimensionality reduction techniques, we chose instead to use *R*_*g*_ and *r*_*e*2*e*_ because they provide intuitive interpretability and clearly capture global conformational changes such as *α*-helical structure, extended structure, and the stability of the initial *β*-hairpin fold versus extended or flexible states. In many cases, the top collective variables extracted from dimensionality reduction techniques applied to peptide and short polymer simulations are in any case closely related to the shape parameters ^47–49^. For each peptide variant, data from 10 independent simulation replicas over the final 400 ns of simulations were combined. The radius of gyration was calculated using MD-Traj ^45^, and the end-to-end distance was determined by measuring the distance between the C_*α*_ atoms of the N-terminal and C-terminal residues, and the free energy surfaces were generated using PyEMMA ^50^(`plot-free-energy` module).

#### 2.4.5 Hydrogen Bonding

Hydrogen bond (hbond) analyses were performed using the `HydrogenBondAnalysis` module in MDAnalysis ^51–53^ to evaluate both intramolecular peptide-peptide and intermolecular peptide–lipopolysaccharide (LPS) interactions. For intramolecular peptide calculations, hydrogen bonds were calculated between peptide atoms within the single peptide chain, whereas peptide–LPS evaluations tracked H-bonds between peptide atoms and non-protein LPS atoms. A standard distance cutoff of 3.5 A and a donor-hydrogen-acceptor angle cutoff of 150 were applied to both analyses ^22^. The results were calculated across the final 400 ns of production trajectories from 10 independent replicas. Per-residue occupancy percentages were computed as the fraction of trajectory frames in which that residue participates in a valid hydrogen bond, and final reported values represent the mean and standard error of the mean across all replicas.

#### 2.4.6 Construction of the potential of mean force

The potential of mean force (PMF) binding profiles were constructed by combining the biased histograms from all 30 umbrella sampling windows using the Weighted Histogram Analysis Method (WHAM) ^54^ via the standard GROMACS command-line tool. Statistical uncertainties and profile convergence across the three independent replicas were assessed to determine the final binding free energy profiles.

## 3 Results and Discussion

We perform molecular dynamics simulations of GL13K and its two variants, GL13KS9A and GL13KS9T, to understand their interactions with LPS, with a specific focus on the role of the serine residue at position nine and the effects of its mutations on the peptide shape and peptide-LPS interaction.

### 3.1 Initial starting structure does not bias peptide-LPS interactions

Initially, we calculated the center-of-mass (COM) distances between the peptide and the lipid A, core, and O-antigen regions over time for both models. As expected, in the first initial structure Fig. 1-b, the peptide remains localized at the lipid A/core region, while in the other starting structure, Fig. 1-c, the peptide migrates from the O-antigen moiety toward the lipid A/core region, stabilizing within approximately 100 ns ( S1). Next, we performed a more detailed analysis by examining the average center-of-mass (COM) distances between the individual residues of the peptide and the individual residues of the lipid A/core region during the final 200 ns of the simulation after all replicates had migrated to their final positions. Fig. 2 shows the residuewise interaction profile between the peptide and the lipid A/core regions. (Intrapeptide contact maps are shown in S3). Across all systems, the peptides localize near the lipid A/inner core, showing a consistent structure in which the N-terminus extends toward the outer core, while the C-terminus remains anchored near the lipid A region. All three peptides show a similar pattern of residue-residue interactions, though we note that N-terminal end of GL13K shows a tendency to be on average slightly more distant from the LPS than the two variants. In particular, residues 1 and 2 of GL13K remain further than about 1.5 nm from all residues, while residue 1 of GL13KS9A approaches within about 1 nm of two of the phosphorylated acyl chains (residues 4/5), while residues 1 and 2 of GL13KS9T also approach within about 1 nm of residue 4 of LPS.

**Fig. 2.**
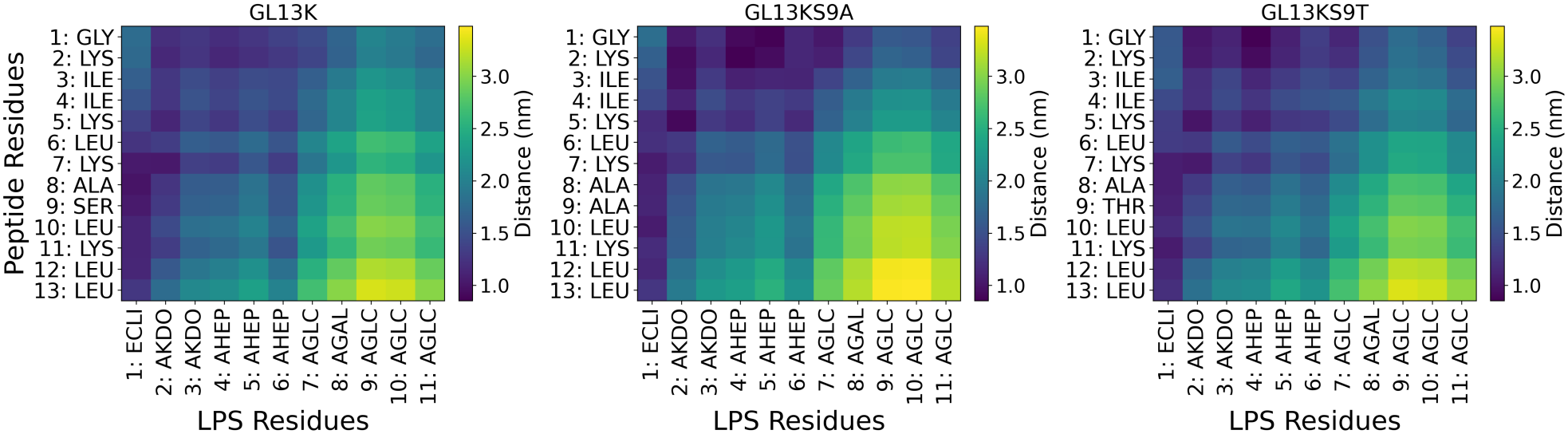
Residue-wise contact map between the peptide and the lipid A/core regions for all peptides, showing the average equilibrium distance (nm) between the centers of mass of peptide residues and (larger) LPS residues. The standard error is plotted in S2.

While all three peptides consistently prefer the lipid A region over the O-antigen and the previous analysis suggests differences between C-terminal and N-terminal positioning across the variants, it does not highlight specific interactions. To address this, we focus on the behavior of the C-terminal and N-terminal residues, and the two closest charged residues to the termini. In Fig. 3, we calculated the average radial distribution functions (RDFs) of the lipid A and core phosphate groups with respect to the N-terminus (GLY01), the adjacent lysine (LYS02), the C-terminus (LEU13), and the nearest lysine to C-terminus (LYS11) for the last 400 ns of simulation using the compute_rdf module in MDTraj ^45^. The RDF profiles show that the C-terminus and LYS11 exclusively interact with the phosphate groups of Lipid A (Fig. 3, third and fourth rows, blue line nearly flat across). The broad radial distribution function of the C-terminus with the Lipid A phosphate (Fig. 3, fourth row, red) shows non-specific interactions between the C-terminus and Lipid A, suggesting that they do not bind but merely tend to remain anchored within about 1-1.5 nm distant from each other.

**Fig. 3.**
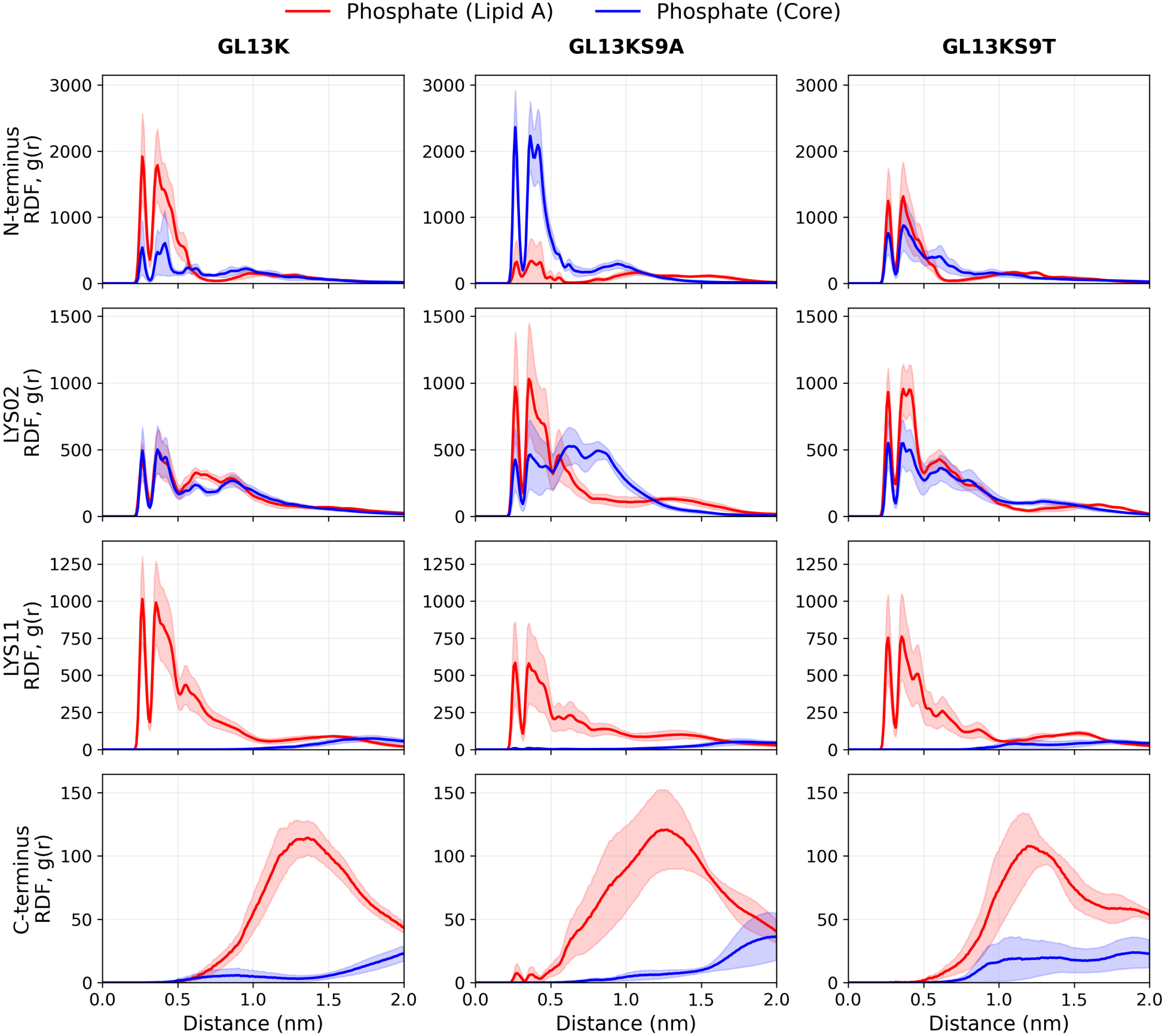
Average radial distribution functions (RDFs) of the lipid A and core phosphate groups with respect to the N-terminus (GLY01), the adjacent lysine (LYS02), the C-terminus (LEU13), and the nearest lysine (LYS11) for the last 400 ns of simulation for all three peptides across all replicas. The standard error is shown as the shaded area which is calculated across all replicas.

Conversely, the N-terminus and LYS2 interact with both the phosphate groups of Lipid A and with the phosphate groups of the core, but they demonstrate differential interactions according to sequence. Specifically, the N-terminus of GL13K preferentially binds with the two phosphates of Lipid A (the two peaks in the RDF, red much greater than blue), while the N-terminus of GL13KS9A preferentially associates with the core phosphate region (blue peaks, no red peaks at all), and GL13KS9T N-terminus interacts with both regions via conformational rearrangement. The LYS02 residue for all three variants interacts with both the phosphates of Lipid A and the core, but the interactions of GL13KS9A and GL13KS9T are stronger between the lysine and the Lipid A phosphate than those of the wild type. In one configuration, it is possible for LYS02 to interact simultaneously with both the Lipid A and core phosphate groups, which have an average distance ranging from about 10 to 15 Å: the charged *ε*-amino group of LYS02 can extend to interact with Lipid A phosphates, while the backbone *α*-carbon remains positioned further away (at approximately 5-6Å). This position is responsible for the appearance of the third and fourth peaks (between 5-10Å) in the LYS02 RDF, implying switching between this configuration and the configuration in which the *α*-carbon is closer to the core phosphates (the first and second peaks, at 5Å). Additionally, due to the strong interaction between the GL13KS9A N-terminus and the core region, alongside the higher peak between LYS02 and the lipid A phosphate groups (red line), we propose that the N-terminal-adjacent residues of GL13KS9A become sandwiched between the lipid A and core regions. In contrast, wild-type GL13K does not exhibit this behavior. Instead, consistent with the heightened interaction of the N-terminus withe Lipid-A phosphates along with lower but present LYS02 interaction peaks with the Lipid-A phosphates, we observe a bend near the N-terminus (ILE03, cf. also Section **??**, Fig. 6) that redirects the remaining lysine residues back towards Lipid-A but at a slightly further distance on average. Furthermore, the non-exclusive interactions of the N-terminus and LYS02 with both the core and Lipid A moieties provide the structural flexibility necessary for the peptide, specifically, the wild-type GL13K, to explore more compact conformational states.

In summary, across all three peptides, Lys11 and the C-terminus show distinct interaction profiles with the lipid A region: the C-terminus shows non-specific interactions, whereas Lys11 displays a higher, more localized RDF peak. On the other hand, the N-terminus shows considerably greater conformational flexibility and a different binding behavior. Specifically, while the N-terminus of wild-type GL13K preferentially associates with the lipid A region, the N-terminus of GL13KS9A almost exclusively binds to the core region.

In summary, across all three peptides, Lys11 and the C-terminus show distinct interaction profiles with the lipid A region: the C-terminus shows non-specific interactions, whereas Lys11 displays a higher, more localized RDF peak. On the other hand, the N-terminus shows considerably greater conformational flexibility and a different binding behavior. Specifically, while the N-terminus of wild-type GL13K preferentially associates with the lipid A region, the N-terminus of GL13KS9A exclusively targets the core region.

### 3.2 LYS07 Anchors the Peptide to Lipid-A through H-bonding

To further evaluate residue-level contributions to complex stability, we calculated the residue-based hydrogen-bonding occupancy between the peptides and LPS over the final 400 ns of the simulation trajectories. As shown in Fig. 4, lysine residues across all three peptides form extensive hydrogen bonds with LPS, with Lipid-A exhibiting the highest occupancy by a wide margin. Specifically, LYS7, positioned centrally in the peptide sequence, displays the highest hydrogen-bonding occupancy ( 82%*±*3%) in GL13K, compared to 58%*±*9% and 65%*±*18% in GL13KS9A and GL13KS9T, respectively, while in GL13KS9A and GL13KS9T, the highest hydrogen-bonder is LYS5 ( 65.8%*±*8.8% and 64.9%*±*15.5%). Additionally, GL13K and GL13KS9T display a fairly substantial amount of hydrogen-bonding between the N-terminus and Lipid A, while GL13KS9A does not (< 7%), with its N-terminus hydrogen-bonding primarily with the inner core phosphates instead, commensurate with the RDF. Similarly, LYS2 in all three variants hydrogen bonds with Lipid A and to a lesser extent to the inner core phosphate region. Surprisingly, residues at position 9 fail to form substantial hydrogen bonds with LPS. For Ala9 in GL13KS9A, interactions are of course restricted exclusively to the backbone due to the nonpolar methyl side chain, but both SER9 and THR9 in GL13K and GL13KS9T exhibit lower occupancies of only 10%*±*8% and 15%*±*11%, respectively.

**Fig. 4.**
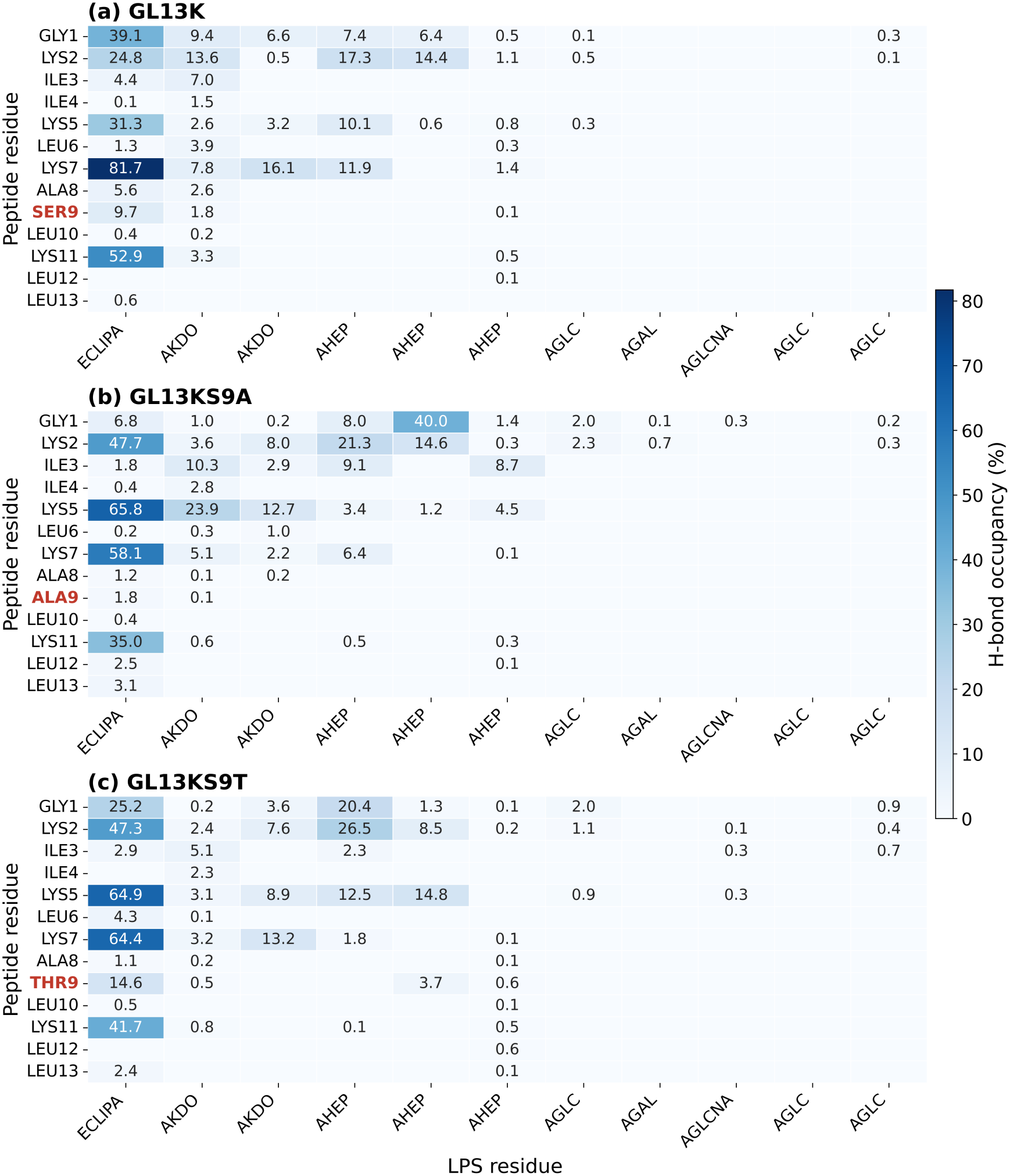
Peptide-LPS h-bond occupancy for Lipid A and the core region, calculated for three peptides across 5 replicas for the last 400 ns. The error values plot is presented in S7.

Aside from the Lipid A moiety, hydrogen bonding between the core region and the peptides occurs primarily at AHEP residues, where the phosphate groups are located. While electrostatic interactions are expected to dictate the association between charged peptide residues and LPS, the radial distribution function results (RDF, Figure 3) and h-bonding results reveal a notable and perhaps more subtle shift: a non-charged SER to ALA mutation in GL13KS9A redirects both the N-terminal GLY and its hydrogen bonds from Lipid A to the core region, while maintaining or strengthening LYS2 interactions with both Lipid A and the inner core. Furthermore, as will be demonstrated in subsequent sections, this single mutation significantly restricts both the conformational flexibility of GL13KS9A and its overall binding affinity for LPS.

### 3.3 Alanine Substitution Creates a Two-State System

To investigate the changes in the conformational landscape of the bound peptide between the three variants, we compute two-dimensional representative free energy surfaces of GL13K and its variants as a function of the radius of gyration, *R*_*g*_, and end-to-end distance *r*_*e*2*e*_. These free energy surfaces (FES) are constructed in a way to support two main objectives: a high level of interpretability and second, capturing desired slow processes. It is possible to use more sophisticated collective variables (CV) than *R*_*g*_ and *r*_*e*2*e*_, however, we argue that, in this project, interpretability is more important than the possibility of finding the slowest process using complex, less-intuitive CVs.

Fig. 5 shows that GL13K explores more states and is more flexible than the other two variants. Specifically, the energy barriers connecting its conformational basins are lower for GL13K than for the variants, which can get as high as 2 to 3 *k*_*B*_*T* and the metastable states are much more spread out (substantially wider purple area in the leftmost panel). (It is important to mention that the barrier energies are certainly within the biological energy fluctuation rate of ~ 5*k*_*B*_*T* –observed landscape changes are comparatively subtle.) Additionally, GL13KS9T has a more compact and constrained structure than the other two peptides, as can be seen by the reduced space it samples in *R*_*g*_: while the other two reach *R*_*g*_ values of 1.3 nm, GL13KS9T barely fluctuates beyond 1.2.

**Fig. 5.**
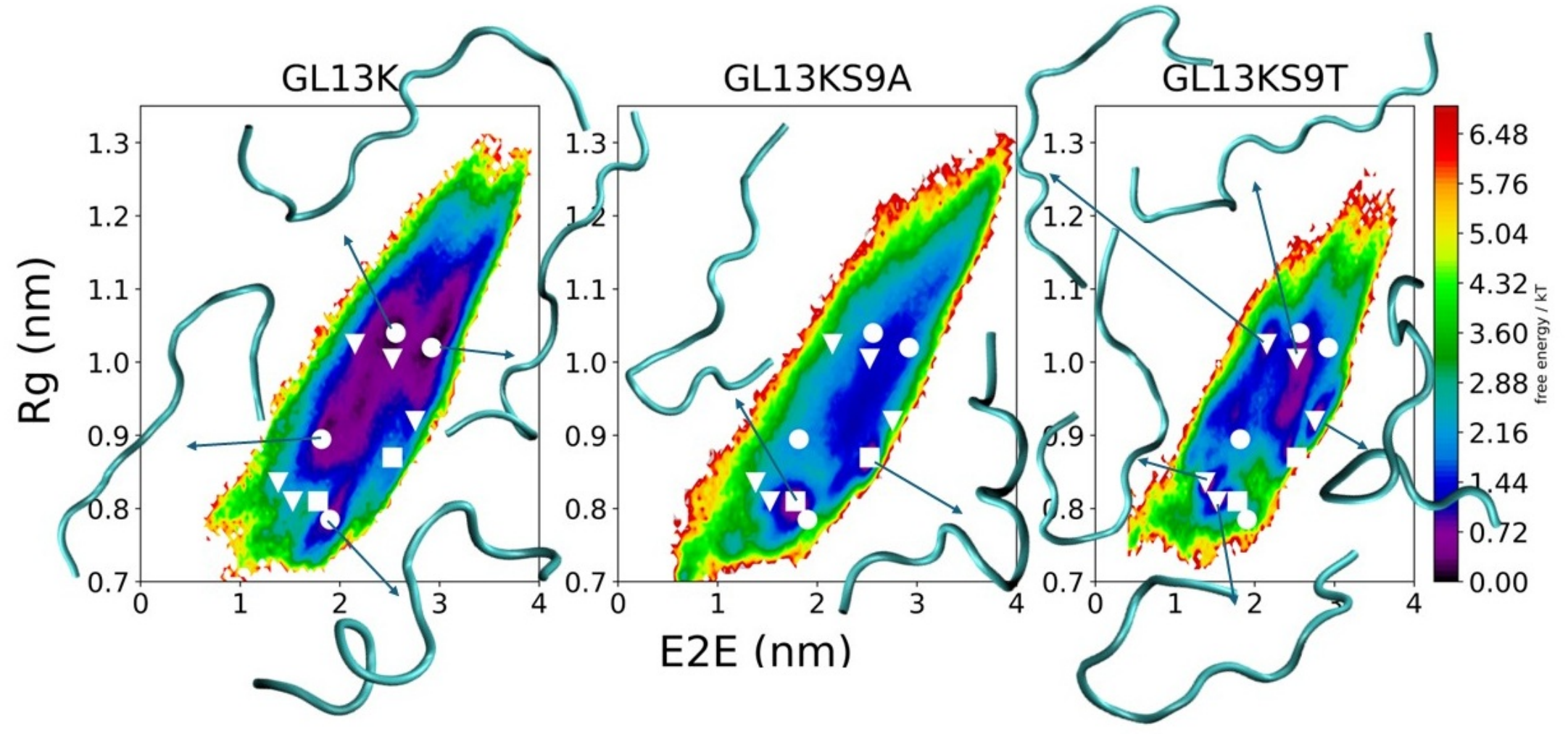
Free energy surface of three peptides constructed by *R*_*g*_ and *r*_*e*2*e*_ variables over the last 400 ns. Snapshots for basins are added to the plot. The basins of GL13K, GL13KS9A and GL13KS9T are represented by white circle, square, and triangle, respectively.

**Fig. 6.**
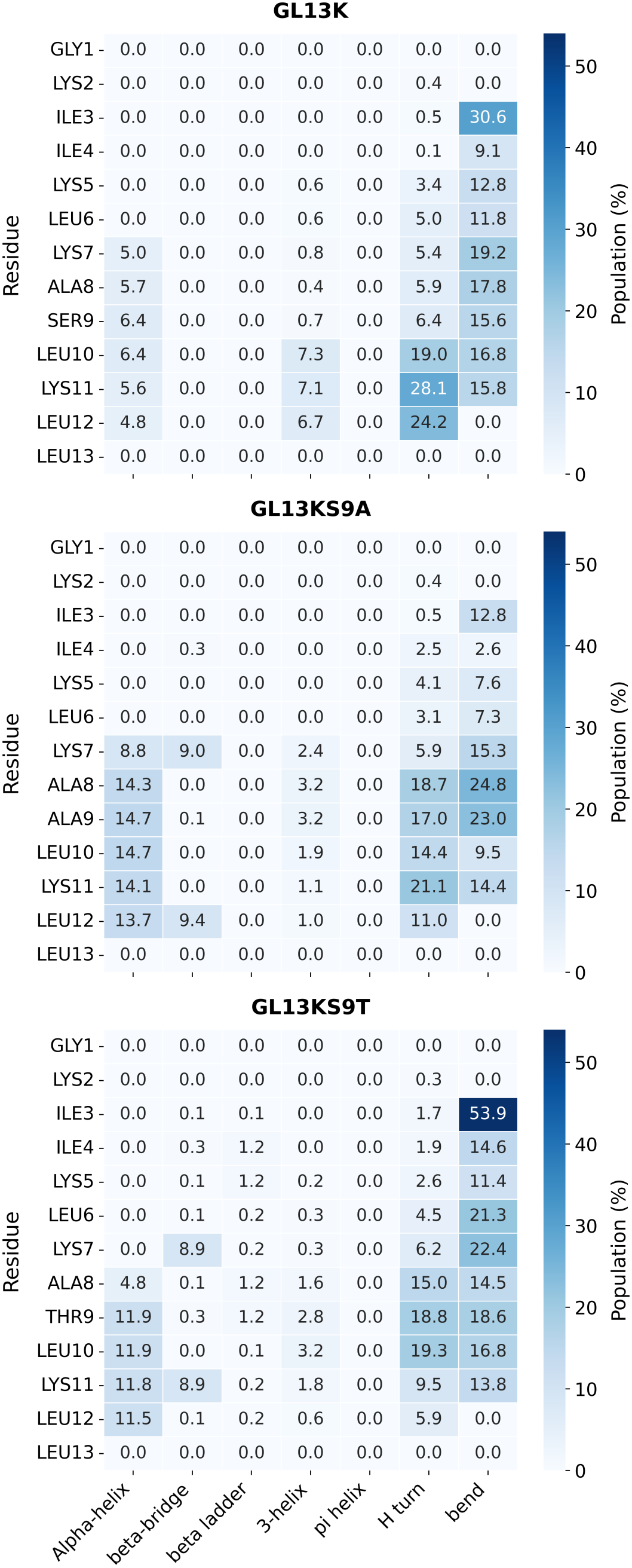
Secondary structure per residue of the three peptides over the last 400 ns, excluding loops content. The error plot is provided in S5

The observations of the local minima on the FES plots show no direct overlap among the three peptides (minima for GL13K, GL13KS9A, and GL13KS9T are indicated by white circles, squares, and triangles, respectively.) Nonetheless, the shapes associated with the minima as well as their proximity and steepness give us important information about the conformational landscape. Of the three, GL13KS9A is the most restricted in terms of favorable metastable states, essentially forming a two-state system with collapsed minima (*R*_*g*_ < 0.9), corresponding to kinked conformations that still have relatively higher end-to-end distance. GL13K has a relative nearby minimum which is separated by a small barrier from the rest of its metastable states (at around *R*_*g*_ *≈* 0.78 nm, *r*_*e*2*e*_ *≈* 2 nm; however, it also occupies substantially more low free-energy conformational space as noted above. GL13KS9A forms more of a multi-state system with about five distinct metastable states, three of which are close to overlapping with the broader metastable surface of GL13K, two of which are folded over with a lower end to end distance than GL13K or GL13KS9A’s minima, though all three variants sample the locations of one another’s minima to an extent. Overall, the SER09 to ALA09 mutation in GL13KS9A significantly reduces conformational flexibility in the LPS-bound state. For GL13KS9T, the mutation from SER09 to THR09 not only decreases conformational flexibility compared to GL13K but also restricts the range of the radius of gyration (*R*_*g*_). We propose that this *R*_*g*_ limitation in GL13KS9T arises from the extra methyl group on the THR09 side chain, which restricts the backbone movement, resulting in lower *R*_*g*_ values relative to the other two peptides.

To complement the FES analysis, we examined whether the observed conformational basins correspond to specific secondary structures by mapping the helical and *β*-strand fractions onto the scatter plots (S4). We note that both basins of the GL13KS9A system occur in a region of heightened helical content, with certain snapshots that occupy this region possessing a helical fraction of about 40-50%. Thus, GL13KS9A demonstrates partial helical folding which stabilizes its two most stable states. Conversely, the most metastable regions of the other two sequences are less structured (containing snapshots that are mostly turns, bends, and coils.)

To further assess secondary structural features of the peptide, we plotted the average secondary structure per residue in Fig. 6. The secondary structure population heatmaps reveal how point mutations at position 9 alter the peptide’s conformational behavior when bound to a single LPS molecule. As indicated in the scatterplots (S4), the wild-type GL13K peptide exists as a disordered structure (<7% alpha-helical content per residue), relying on turns and bends with the most notably an H-turn propensity at LEU10, LYS11, and LEU12 and bends across LYS7 to LEU10, to facilitate the binding of a single peptide to LPS. Also, it is important to mention that a 30.6%*±*6.3% bend is observed at ILE3 residue. Introducing the S9A mutation does indeed introduce more structural propensity, with a small amount of alpha-helical conformation observed across residues 7–12 ( 13–15%), due to alanine’s high intrinsic helix propensity. In contrast, the S9T mutation stabilizes the bend at ILE3. This likely occurs because of the introduction of a bulkier, beta-branched threonine residue that heavily restricts local backbone dihedral flexibility near position 9. When cationic residues like LYS2, LYS5, LYS7, and LYS11 form strong electrostatic interactions with the negatively charged LPS phosphate groups, the peptide becomes rigidly anchored to the LPS, and it is unable to smoothly distribute conformational changes due to the restrained threonine backbone. This results in inducing a compensatory bend near the flexible N-terminus that rises the bending population at ILE3 to a dominant 53.9%*±*8.3%

### 3.4 Shape parameters underscore compactness of GL13KS9T

While GL13KS9T shows a smaller *R*_*g*_ range than the other two variants, it remains unclear whether this behavior arises from true 3D structural compactness or merely an asymmetric mass distribution off the main axis. To address this, Fig. 7 presents the radius of gyration (*R*_*g*_), asphericity (*a*_*sph*_), and acylindricity (*a*_*cyl*_) for each peptide, with standard errors calculated from 10 replicas over the final 400 ns of the simulations. The results show that GL13KS9T has a lower asphericity than GL13K, which, on average, corresponds to a smaller radius of gyration. More importantly, the acylindricity of GL13KS9T overlaps with the other two peptides, ruling out an asymmetric mass distribution off the main axis and confirming that this variant is genuinely more compact overall.

**Fig. 7.**
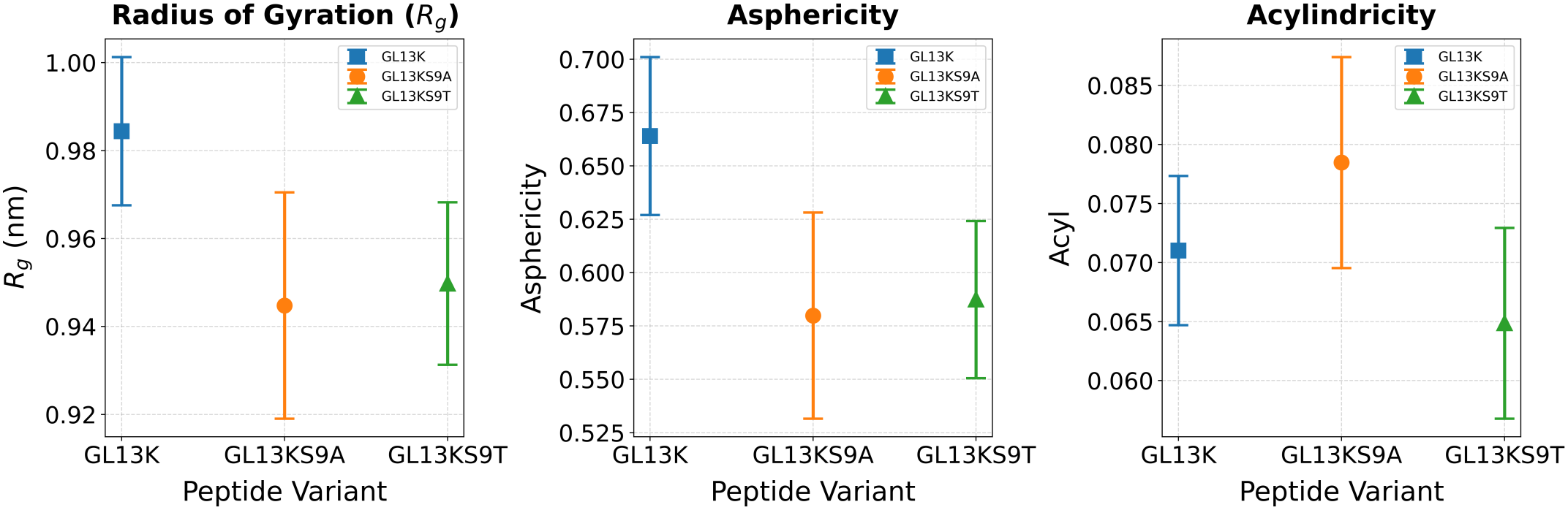
Average shape parameters of GL13K, GL13KS9A and GL13KS9T from left to right: radius of gyration (*R*_*g*_), asphericity (*a*_*sph*_), and acylindricity (*a*_*cyl*_). The error bars indicate the standard error in the means of the ten different replicas.

### 3.5 The Ninth Residue Governs Intrapeptide Hydrogen Bonding

The remaining question is how the substitution of SER09 affects peptide behavior at the atomistic level, particularly as it does not appear to strongly hydrogen-bond with LPS. To investigate this, we calculated the hydrogen-bond occupancy between nonadjacent peptide residues, as shown in Fig. 8. In GL13K, SER09 participates in several prominent hydrogen-bonding interactions, particularly with LYS11 (34.7% *±*5.8%), LEU12 (19.6%*±*7.2%), and LEU13 (15.7%*±*5.4%), suggesting that SER09 contributes substantially to the intrapeptide hydrogen-bond network at C-terminus. Substitution with alanine considerably reduces these interactions, with the occupancy of the LYS11–ALA09 pair decreasing to 12.2%*±*3.1%. In contrast, the threonine substitution largely retains the interaction with LYS11, showing an occupancy of 36.6%*±*13.5%, while the LEU12–THR09 interaction reaches a low 7.0%*±*4.7%. Overall, these observations indicate that SER09 substitution alters the intrapeptide hydrogen-bonding network in a mutation-dependent manner, with threonine retaining some of the prominent interactions observed in GL13K, whereas alanine is associated with a more redistribution of these interactions among near residue to ALA09 while losing LYS11,LEU12 and LEU13.

**Fig. 8.**
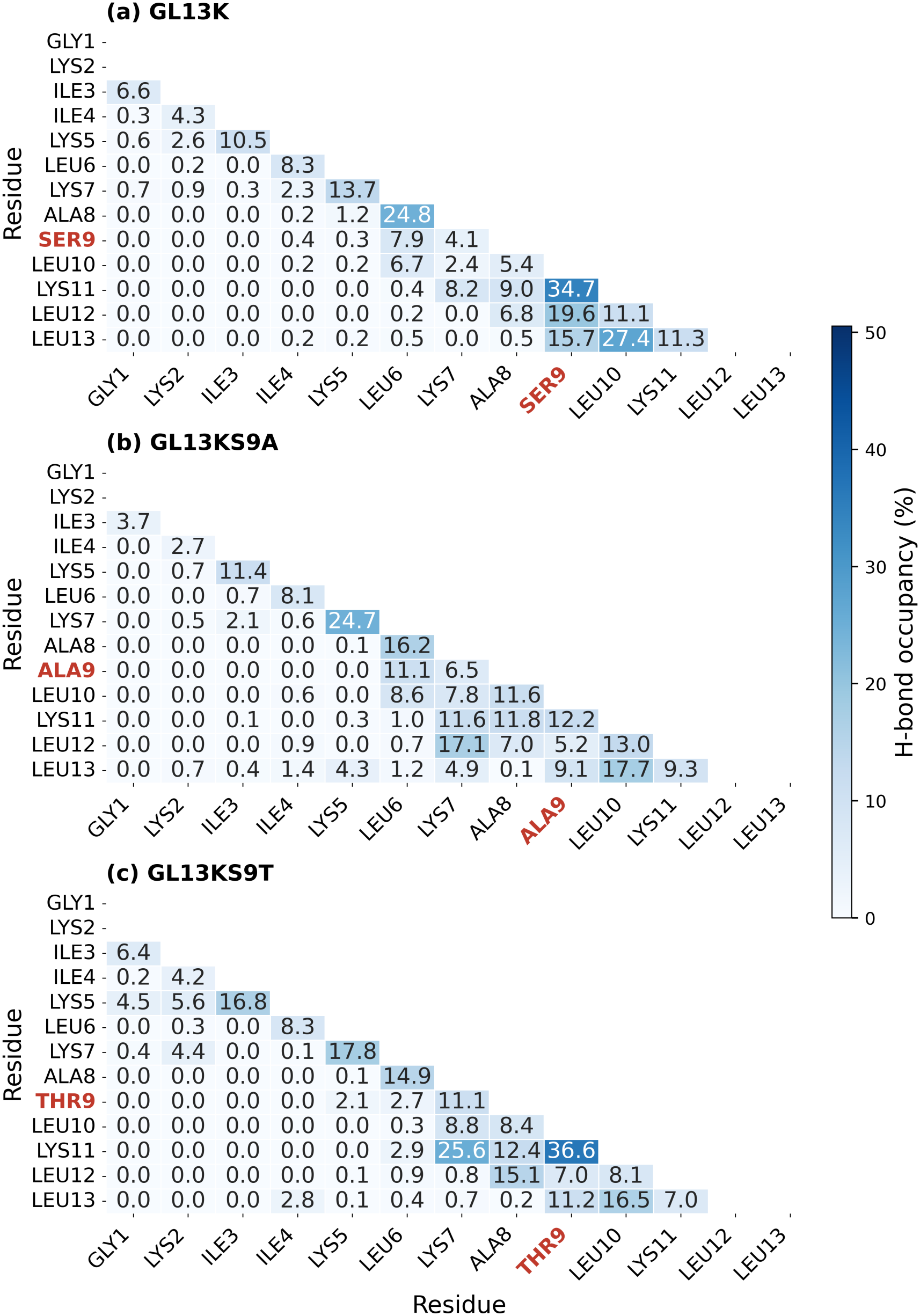
Peptide-peptide h-bond occupancy for non-adjacent residues, calculated for three peptides across 10 replicas for the last 400 ns. The error values plot is presented in S6.

### 3.6 Serine Mutations Reduce the Average Energy Barrier between LPS-Bound and Unbound States

In order to estimate the binding of GL13K and its variants to a single LPS molecule, we performed biased simulations, from which we computed the potential of mean force (PMF), plotted in Fig. 9. For the wild-type GL13K, the free energy rises steeply up to around 3.0 nm before reaching a plateau, resulting in the highest free energy change upon unbinding among all three variants with an average height of 32.1 *±* 3.3 *k*_*B*_*T*, though with notable replica variation (ranging from 27 *k*_*B*_*T* up to 40 *k*_*B*_*T*). The GL13KS9A mutant drastically lowers this unbinding free energy barrier down to an average of 18.1 *±* 1.1 *k*_*B*_*T*, displaying a convergence across all three replicas and a much smoother and lower energy landscape across the reaction coordinate. Intermediate to both is the GL13KS9T mutant, which shows an average barrier height of 25.7 *±* 3.3 *k*_*B*_*T*, rising moderately up to 3.0 nm before leveling off into a plateau near 4.5–5.0 nm. Taken together, the mutation at position 9 significantly alters the binding free energy with LPS, where replacing serine with alanine weakens the overall interaction strength the most, whereas replacing it with threonine retains a moderately strong binding free energy change.

**Fig. 9.**
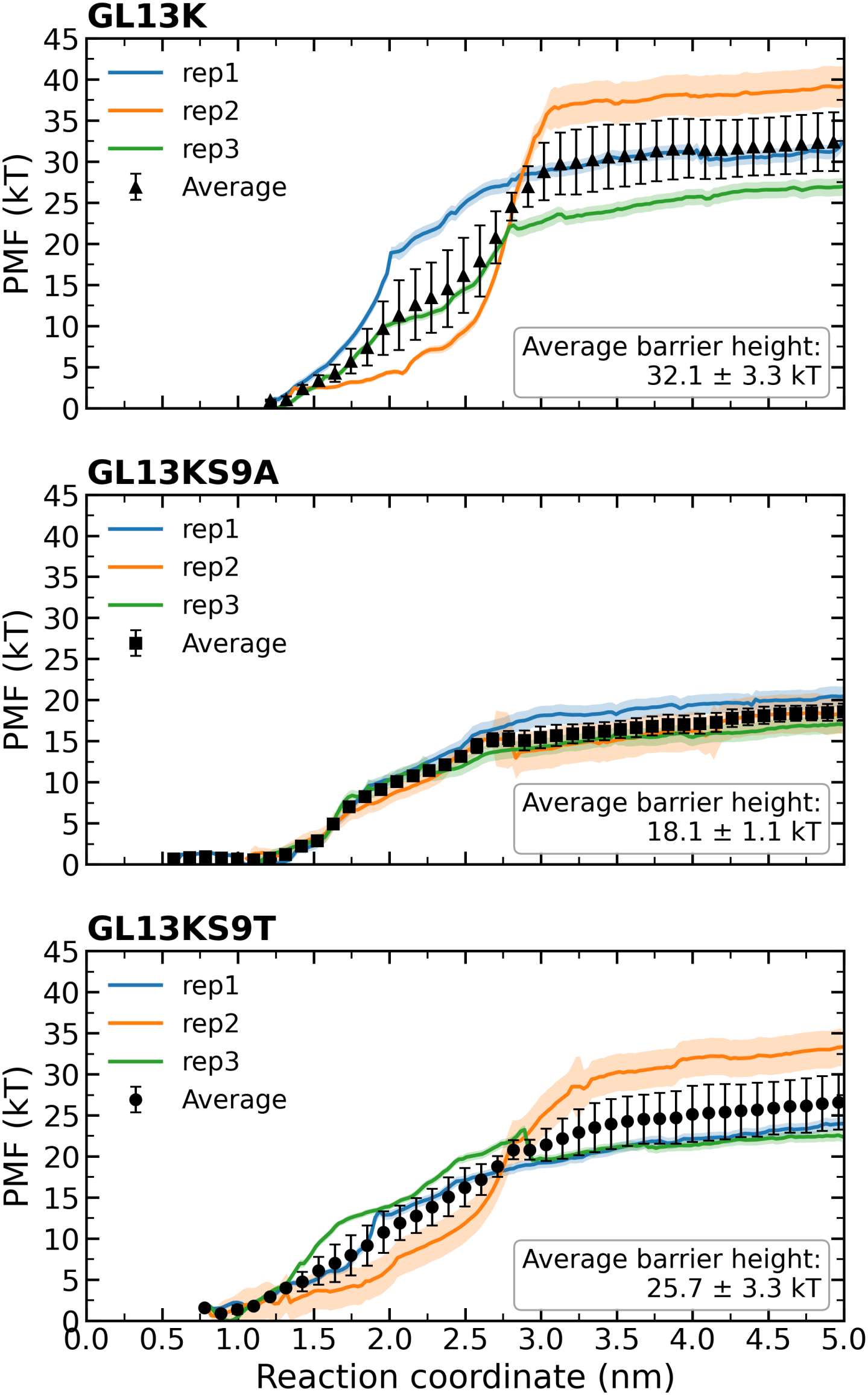
Potentials of Mean Force (PMF) profiles for peptide binding. Free energy profiles along the reaction coordinate for the binding of GL13K, GL13KS9A, and GL13KS9T from an LPS molecule, calculated via umbrella sampling and WHAM at 300 K. Individual independent replicas (three replicas) are shown as colored lines with shaded standard deviation envelopes derived from 300 Bayesian bootstrap iterations (-bsprof). Black data points represent the average PMF profile with standard error bars calculated from the bootstrap distribution (-bsres). Inset: Average barrier heights for each system.

The observed variability among replicas for each peptide is due to a strong dependence on the initial binding pose and, more crucially, the relative orientation of the peptide with respect to the LPS molecule. The direct electrostatic interaction of charged N-terminus and LYS02 with lipid A phosphate group results in the highest barrier energy, an interaction which is greatly reduced in probability in the alanine variant (cf. Fig. 3). In Fig. 10, we show example snapshots of the initial structures of three peptides with respect to LPS, before the umbrella-sampling equilibration step (panel a) and after (panel b). All the peptides are placed almost identically with respect to the lipid A region in the beginning. Only in GL13KS9A, the orientation of the peptide is much more towards the core region, while equilibration of the other two peptides leaves them closer to the Lipid-A region. It is only after equilibration that differences in the peptide orientation with respect to LPS emerge, and while this does lead to variability resulting from initial random velocities of each replicate, it shows that the observed differences in PMF are not due to bias in the initial peptide placement but rather to their differing preferential conformations, as is also supported by the unbiased atomistic simulations.

**Fig. 10.**
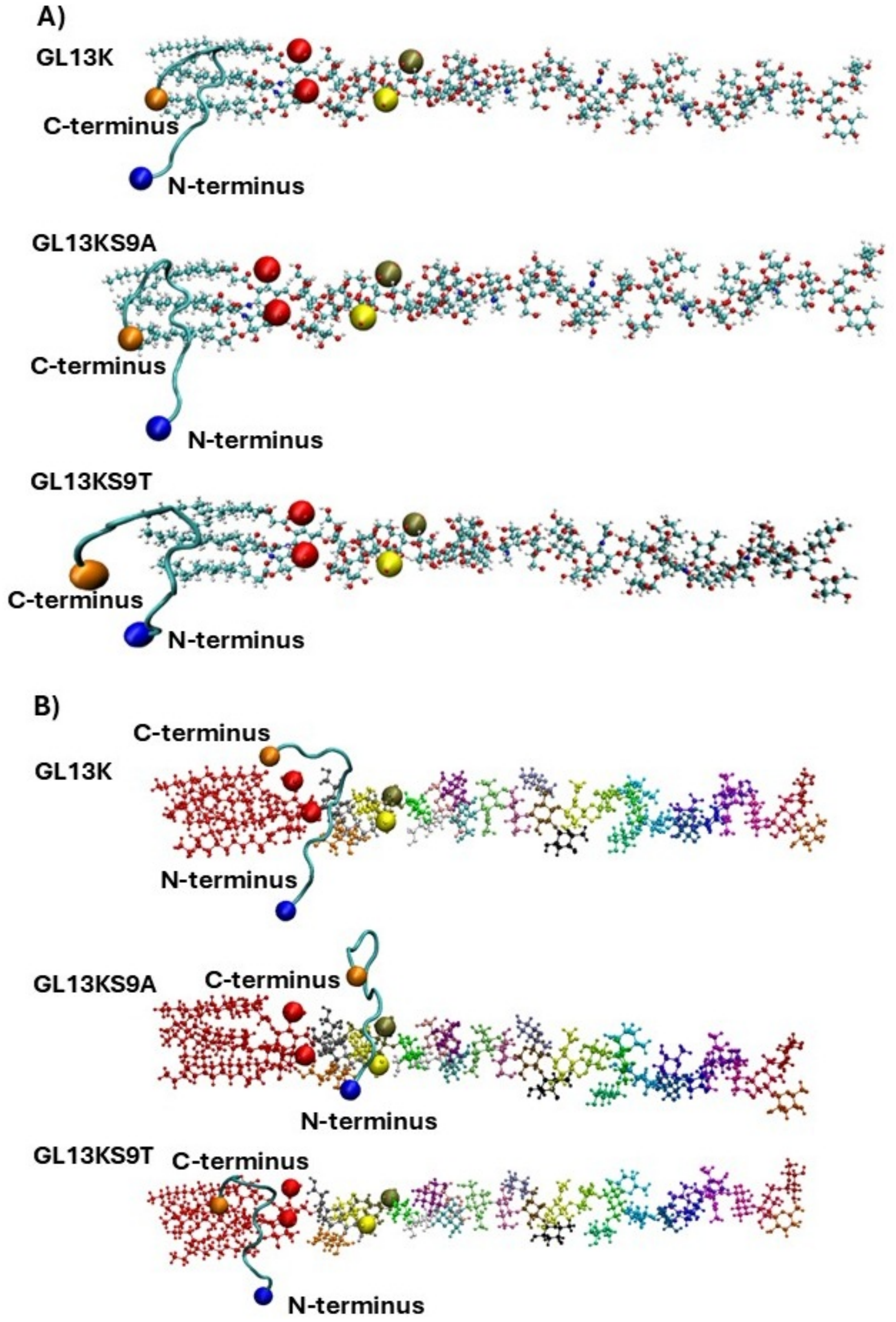
The snapshots of the peptide-LPS complex for all three peptides, A) before the umbrella-sampling equilibrium step (initial structure) B) the first umbrella-sampling window (minimum reaction coordiante value). The N-terminus, the C-terminus and lipid A phosphate groups are shown in blue, orange, and red vdw beads.

## Conclusions

We performed the first *in silico* investigation of a single peptide–LPS interaction, specifically focusing on the conformational and thermodynamic impacts of mutating a single, experimentally-relevant residue. We investigated the overall conformational behavior of the peptide, including the distinct orientational preferences of its N and C-termini during LPS binding and how these affected binding affinities. Crucially, our computational findings closely align with previous experimental observations: the S9A mutation reduces the free energy of binding compared to the wild-type peptide, making it a less favorable process. We are able to identify the likely molecular-level underpinnings of this process: mutation of the ninth residue to a nonpolar alanine disrupts the C-terminal hydrogen bonding network, which shifts the conformational landscape to more strongly favor a transient helical conformation in the C-terminal region over a hook/bend conformation. This shift in the landscape, in turn, results in suppression of a bend in the N-terminus centered at ILE3, corresponding with altered proximity and altered hydrogen bonding of the N-terminal glycine away from the Lipid-A phosphates and towards the inner core heptose phosphates, an orientational shift that observably decreases the overall potential of mean force and thus presumably the free energy of binding. Conversely, the GL13KS9T variant, although it undergoes some slight barrier suppression, probably due to the steric hindrance of the bulkier threonine residue preventing easier conformational rearrangement, does still have a comparable free energy change upon unbinding to the wild type. This underscores the importance of the polar quality of the ninth residue in stabilizing the relevant binding conformations through modifications of the conformational space rather than through directly mediating LPS binding.

Building upon our earlier work ^22^, we hypothesize that the functional significance of SER09 will become even more important in higher concentration peptide systems, where it can actively facilitate interpeptide hydrogen bonding and aggregation of peptide in vicinity of LPS. Notably, while our prior studies on the single, isolated charged peptide in solution ^22^ and at the membrane interface ^23^ under physiological conditions (pH 7.4) revealed a semi-consistent stable *β*-hairpin basin centered at a radius of gyration (*R*_*g*_) of ~0.7 nm and an end-to-end distance of ~0.6 nm, we do not observe a stable *β* hairpin structure in our simulations. Nevertheless, we do observe a comparatively stable and apparently crucial bend in the C-terminus in the wild type and GL13KS9T variant that disappears in the GL13KS9A variant. To build upon these insights, critical avenues for future work include systematically investigating the concentration-dependent interaction of GL13K and its variants with LPS assemblies. Additionally, extending these simulations to complex Gram-negative outer membrane models will be essential to capture peptide penetration dynamics and cooperative interactions across multiple LPS molecules.

## Author contributions

M.N.H. – Conceptualization, methodology, software, validation, formal analysis, investigation, data curation, writing – original draft, writing – review & editing, visualization; J. M. - Conceptualization, methodology – review & editing, visualization; R.A.M. – Conceptualization, methodology, investigation, resources, formal analysis, writing – original draft, writing – review & editing, supervision, project administration, funding acquisition.

## Data availability

The authors confirm that the datasets containing molecular dynamics trajectories without water, structures, force fields and molecular dynamics input parameter files are publicly available at the following DOI:10.5281/zenodo.24338334. Data including the positions of water molecules are available from the authors upon reasonable request.

## Acknowledgements

This research was supported in part by Discovery Grant #RGPIN-2021-03470 from the National Sciences and Engineering Research Council of Canada. This research was enabled in part by support provided by Calcul Quebec (www.calculquebec.ca) and the Digital Research Alliance of Canada (https://alliancecan.ca). This research was undertaken, in part, thanks to funding from the Canada Research Chairs Program under grant number CRC-2020-00225. The authors thank Dr. Christine DeWolf for useful discussions.

