## supplemental figures for "Investigating the critical role of serine in antimicrobial peptide binding to lipopolysaccharide"

### A Supporting Information

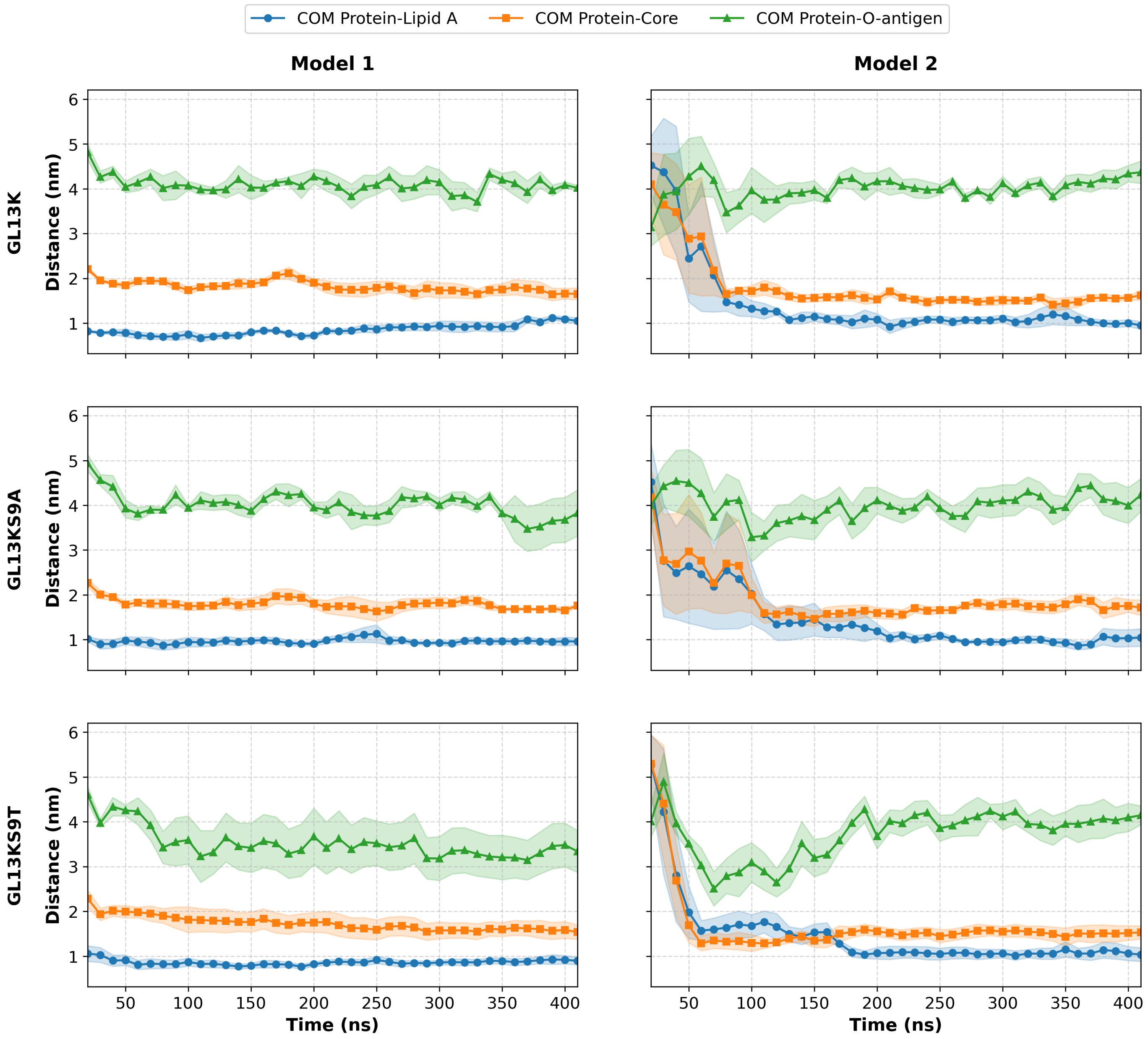


Fig. S1 Average center-of-mass difference distances between peptide and LPS residue wiseddistance over time. The shaded area is calculated as the standard error across 5 replicas.

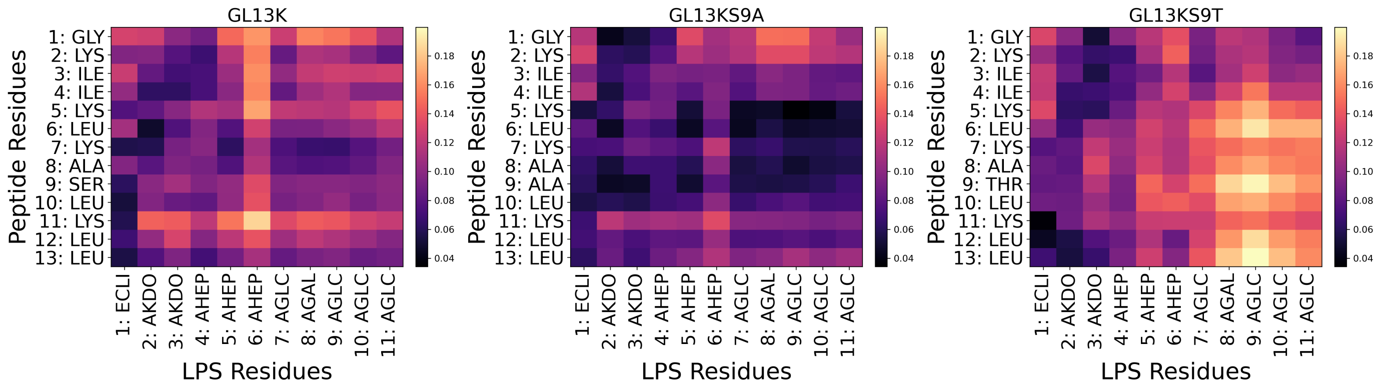


Fig. S2 Standard error of residue-wise interaction profile between the peptide and the lipid A/core regions for all peptides.

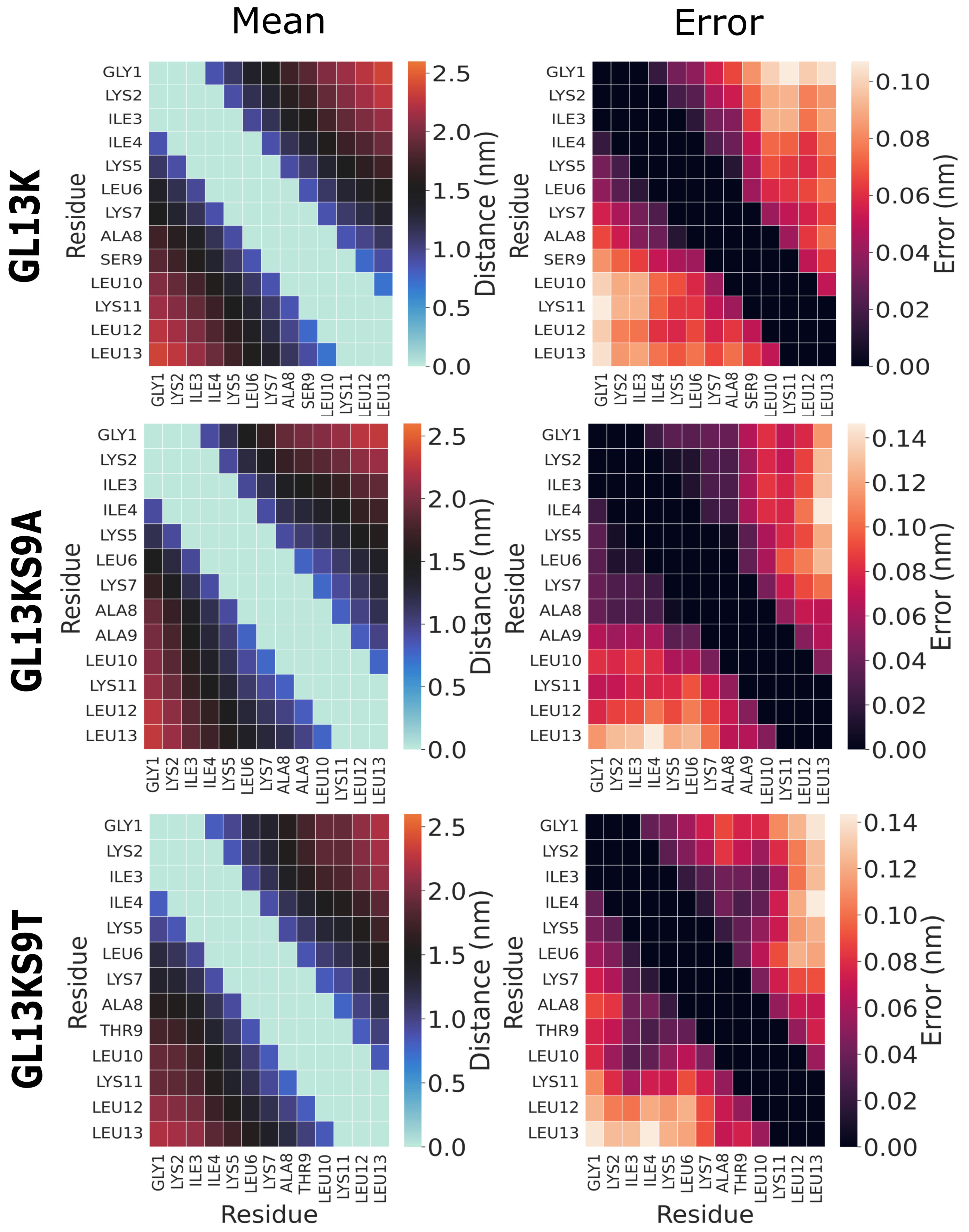


Fig. S3 Peptide-peptide contact map, average over the last 200 ns average and error bars across 10 replicas (model1 and model2).

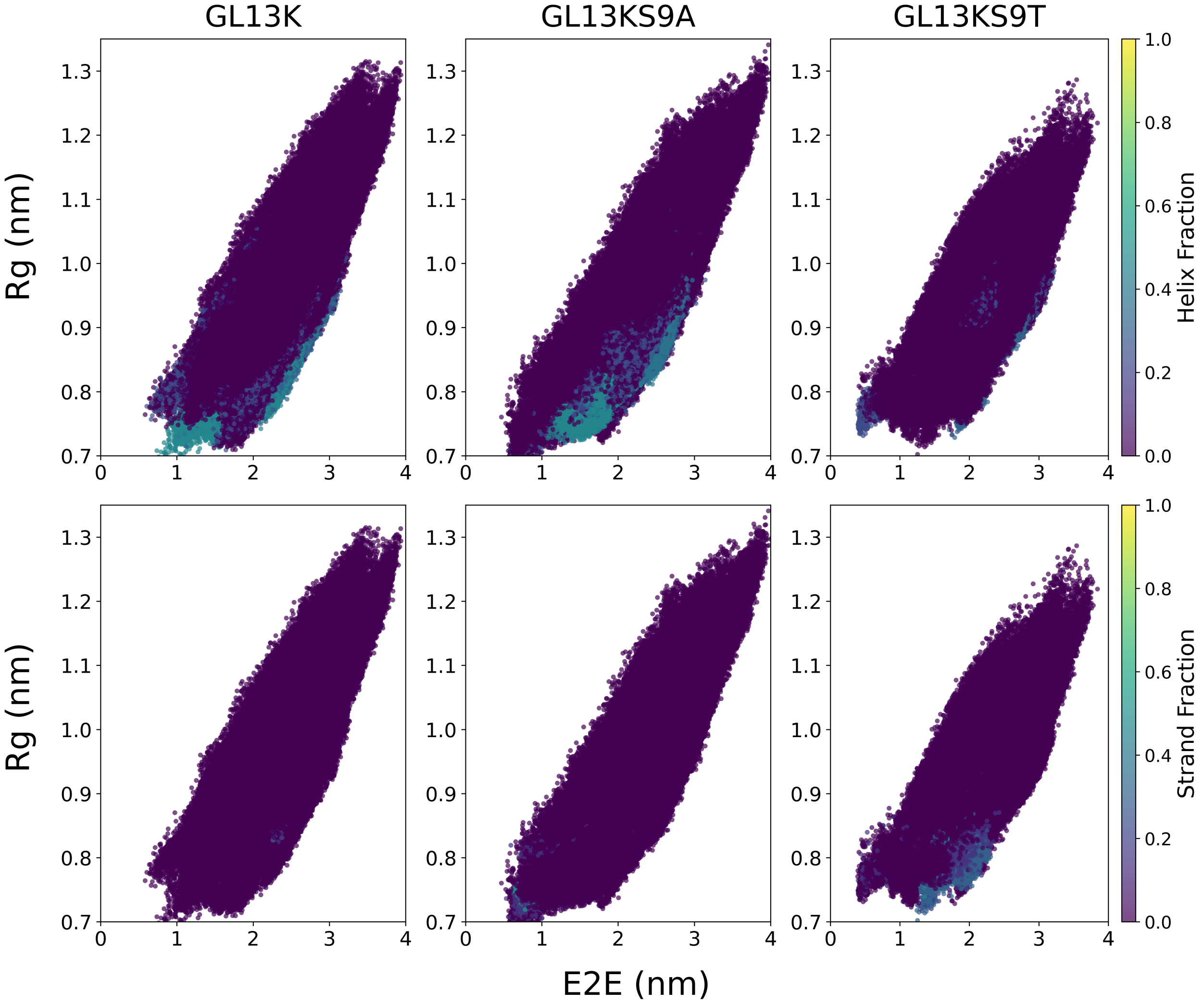


Fig. S4 Scatter plot of the variables used to construct FES, colored by helical (top row) and $\beta$-sheet (bottom row) structural contents, associated with each snapshot of the three peptides over the last 400 ns.

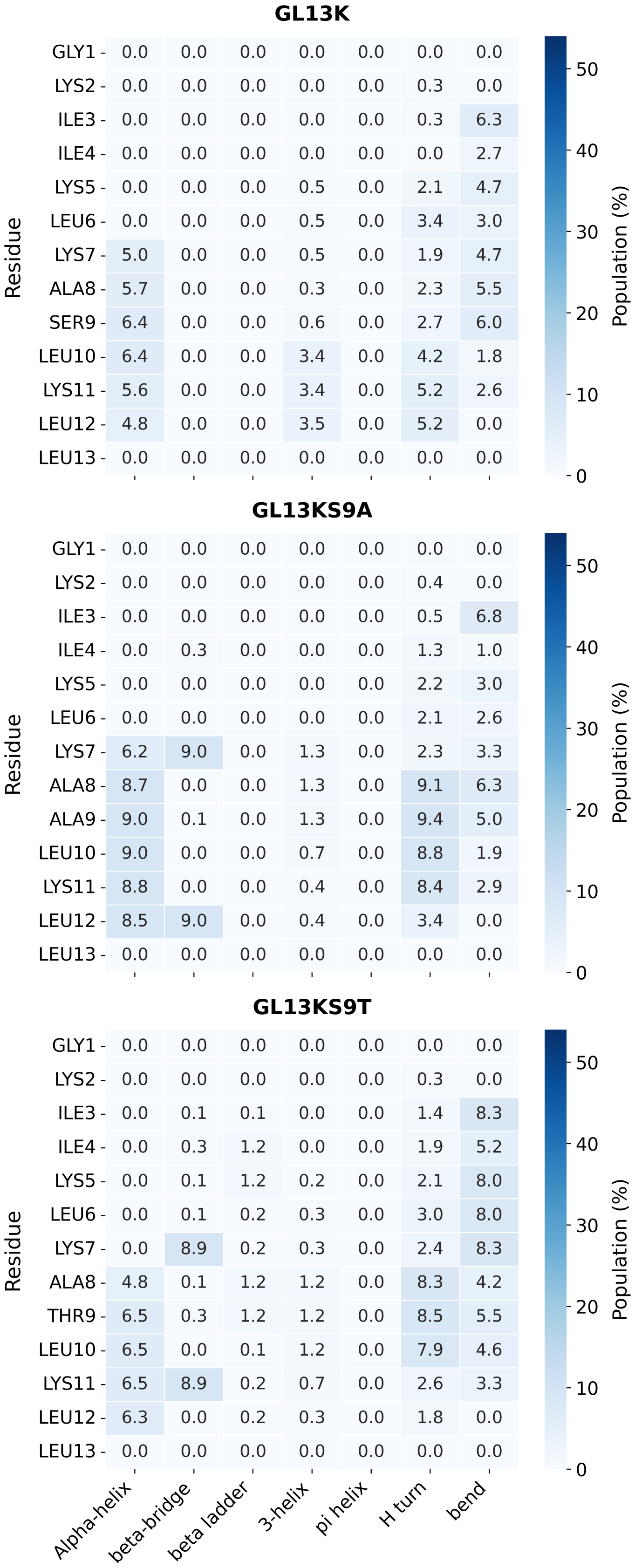


Fig. S5 Standard error secondary structure per residue of the three peptides over the last 400 ns excluding loops content.

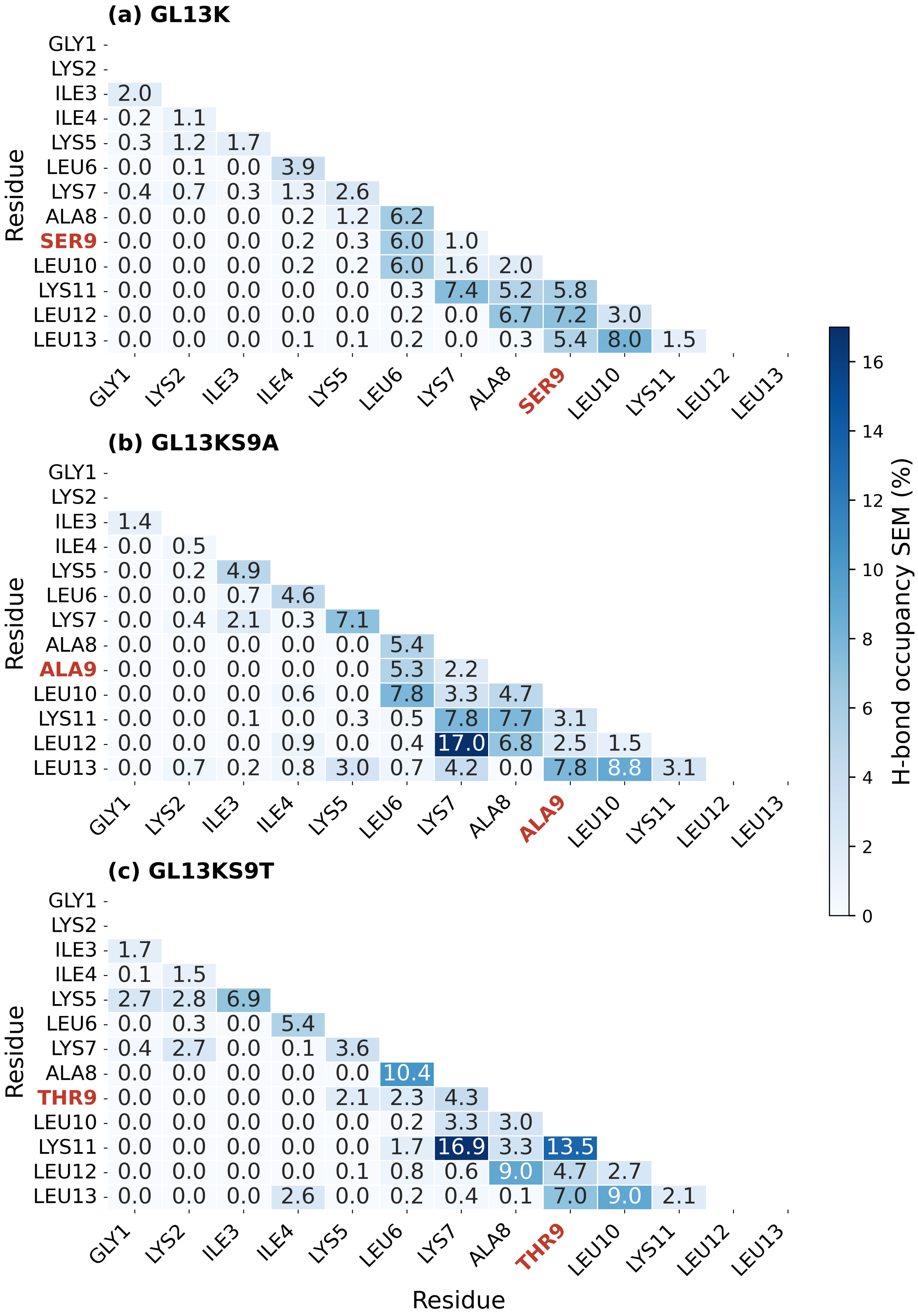


Fig. S6 Peptide-peptide h-bond occupancy error value calculated for three peptides across 10 replicas for the last 400 ns.

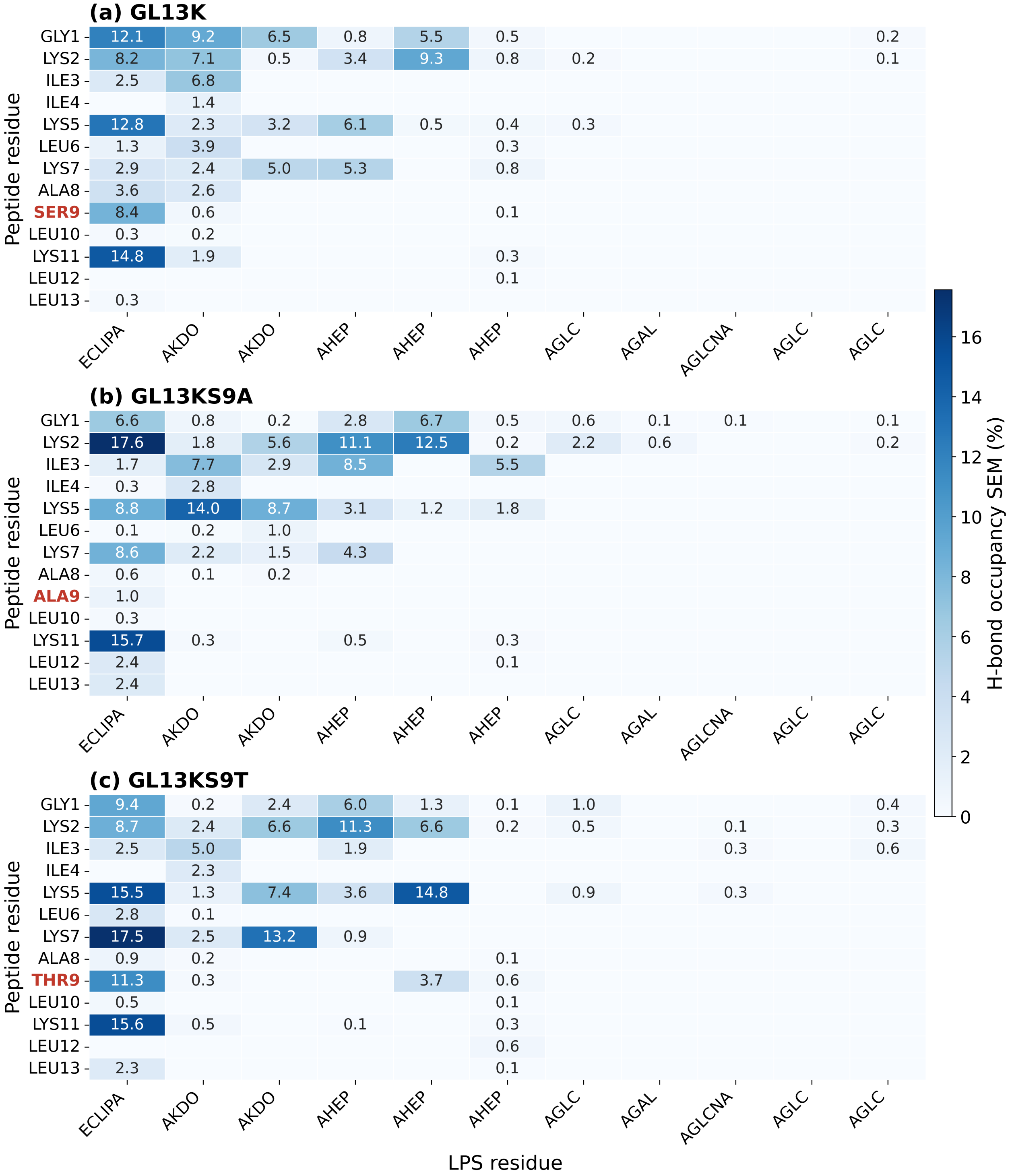


Fig. S7 Peptide-LPS h-bond occupancy error for Lipid A and the core region, calculated for three peptides across 10 replicas for the last 400 ns.
